# Somatic Calcium Signals from Layer II/III Motor Cortex for Continuous Neural Decoding

**DOI:** 10.1101/2023.04.11.536319

**Authors:** Ruixue Wang, Jiawei Han, Xiaoling Hu, Heecheon You, Shaomin Zhang

**Author notes:** Correspondence: Shaomin Zhang. These authors have contributed equally to this work and share first authorship (Ruixue Wang, Jiawei Han).

## Abstract

The latest research shows that calcium signals can provide a new signal source for brain-machine interfaces (BMI). However, it remains a question whether the calcium signals from layer 2/3 motor cortex can be used for continuous neural decoding. And how they are involved in movement coding is also worth investigating. Here we collect the somatic signals in M1 layer 2/3 while mice performed a lever-press task under the one-photon imaging. We first present the potential of somatic calcium signals from layer 2/3 in continuous neural decoding through an improved recurrent neural network. Layer 2/3 neurons exhibit three types of calcium dynamics with distinct spatiotemporal coding patterns involved in the movement. Pre-pressing and pressing neurons enable sparse coding of movement through complementary spatiotemporal information. While post-pressing neurons predict the lever movement most accurately through the calcium dynamics with higher fidelity. These results demonstrate the capability of calcium signals from layer 2/3 neurons as a motor BMI driver and underscore their diversity in motor coding, opening a new avenue for studying the motor cortex and designing optical BMIs.

## 1. Introduction

Population decoding is a powerful way to understand neural content and coding in the BMI system. The signal source is usually electrical signals from the neural activities collected by electrophysiological methods. To date, intracranial microelectrode arrays can monitor activities from hundreds of neurons in non-human primates. The latest Neuropixels probes have been developed to provide approximately 1000 recoding sites on a narrow shank for large-scale neural information recoding[1]. These electrophysiological recording methods have demonstrated great advantages in the neural information source recording. However, they are all spatially sparse and the high-density signals from Neuropixels are not commonly used in population decoding.

Optical imaging methods, as another important method for large-scale neural population recording, have rapidly developed in recent years. Calcium imaging such as two-photon and one-photon imaging methods has been widely used in many studies for learning neural population dynamics[2; 3; 4]. It provides a more comprehensive view of neural activity in a dense, spatially localized, and genetically annotated map[5]. More importantly, it enables us to track the subpopulation of neurons for many weeks or even months[6]. Although the temporal resolution of calcium imaging methods is not as fine as that of electrophysiological techniques. The advantages of calcium imaging methods may help make up for some limitations of current electrophysiological recording methods. The combination of calcium imaging methods and BMI will open up new opportunities for a generation of optical BMIs.

In recent studies, calcium imaging methods, especially two-photon calcium imaging, have been combined with BMI and proved successful[7; 8]. But animals need to keep head-fixed under the traditional two-photon calcium imaging, thus restricting various behavioral tasks. Some differences may be induced by head fix in comparison to free movements as well. While one-photon calcium imaging allows free moving and thus avoids these problems.

In traditional motor BMIs, electrical signals of pyramidal neurons (PNs), typically collected from layer 5b in the primary motor cortex (M1), are served as the signal source of BMI. However, the accessible depth of surface calcium imaging was approximately 600μm, making it challenging to image somatic signals from layer 5b without tissue damage [9]. It is unclear how this tissue damage influences normal neural activities. Therefore, Trautman E *et al*. imaged apical dendritic calcium signals originating from layer 5 output neurons in superficial layers as an agent via two-photon imaging [10]. Masamizu Y *et al*. conducted two-photon calcium imaging in layer 5a at a depth of 503±29μm instead [11]. Actually, there are two intermediate layers upstream of the motor-output layer in M1: layer 2/3 and layer 5a. Layer 2/3 accepts input from the somatosensory cortex [12], which means layer 2/3 neurons directly link somatosensation and control of movements [13]. Previous studies have reported that motor learning caused plasticity in M1 layer 2/3 neurons [14]. The dynamics of layer 2/3 neurons in M1 during a lever-pull task could reflect their high plasticity activities associated with motor primitives and sensory feedback evoked by the lever-pull movement [15]. Theoretically, layer 2/3 neurons integrate sensory and motor information and could function as a signal source of BMI.

In one of our previous studies, we demonstrated that somatic signals from layer 2/3 in the motor cortex were able to decode the discrete movement states [16]. However, whether the temporal resolution of calcium signals is sufficient for continuous neural decoding remains unknown. Masamizu Y *et al*. and Hira R *et al*. used the activities of layer 2/3 neurons in M1 under the two-photon imaging to decode lever trajectory during a lever-pull task [11; 17]. But their two-photon images only provided the highest decoding accuracy about 0.6. Obviously, it is controversial that the calcium signals from layer 2/3 can be used as the electrophysiological signals from layer 5b PNs did in a motor BMI decoder.

In this study, we demonstrated calcium signals from layer 2/3 motor cortex under the one-photon imaging when mice engaged in a lever-press task. We first presented the potential of calcium signals of layer 2/3 neurons in continuous neural decoding through an improved recurrent neural network (RNN). Then we further explored how these layer 2/3 neurons were involved in the movement encoding. There were three types of calcium dynamics displaying distinct spatiotemporal coding patterns. They all specified the lever movement through the ensemble responses. Pre-pressing and pressing neurons predicted the lever trajectories sparsely through complementary spatiotemporal information. Post-pressing neurons enabled the most accurate coding of movement through the calcium dynamics with higher consistency from trial to trial. Our demonstration illustrates new insights for motor cortex investigation and helps to develop a novel optical BMI system.

## 2. Results

### 2.1. One-photon calcium imaging of somatic signals from layer 2/3 motor cortex during a lever-press task

To investigate whether the calcium signals from layer 2/3 neurons were involved in forelimb movement, we trained mice to perform the lever-press task. In each trial, water-restricted mice received a water reward by pressing a lever beyond two set thresholds with their left forelimb during the response time (Figure S1C, Supporting Information). After 10 days of the train, the mice could perform the lever-press task skillfully. A total of 17 session datasets from 6 mice after the train was collected. The mean success rate achieved 85.64 ± 6.50% across all sessions (**Figure 1**A). One-photon imaging was obtained from the left caudal forelimb area (CFA) via UCLA miniscope in a field of view of 700×450 μm (Figure 1B). 144 ± 86 (mean ± SD) ROIs were imaged in each session (a total of 2448 ROIs). ROIs with their corresponding fluorescence were identified after MIN1PIPE processing[18] (Figure 1C, 1D). Different neurons displayed distinct activity patterns. Some neurons mapped well to lever movement while others did not. It is also worth noting that some neurons exhibited a very sparse active pattern (Figure 1D).

**Figure 1.**
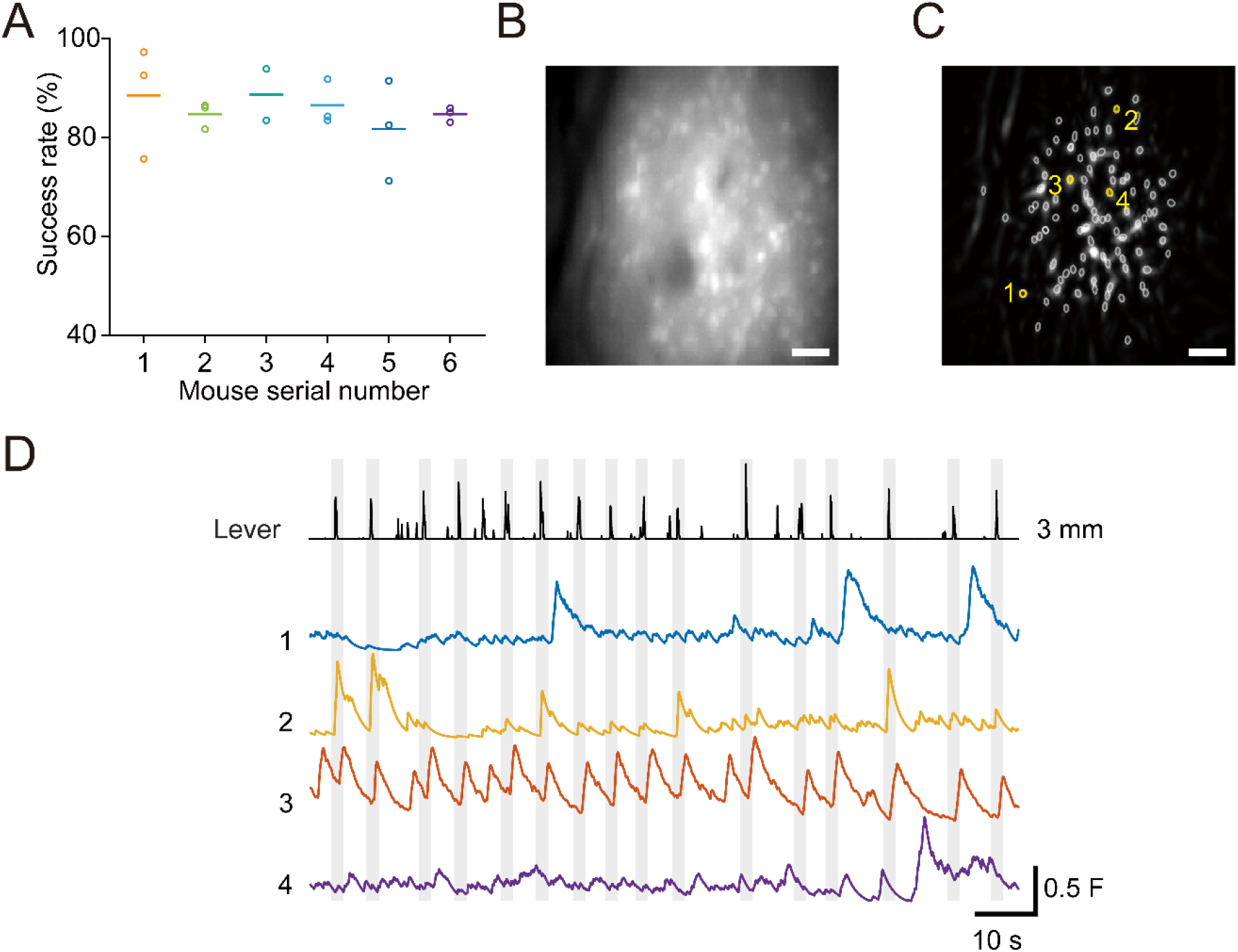
One-photon calcium imaging during a lever-press task. (A) Behavioral performance. The mean success rate for 17 sessions from 6 mice. (B) A representative one-photon image of layer 2/3 neurons expressing Gcamp7f in the left CFA. Scale bar, 50μm. (C) All reconstructed ROIs are outlined within the imaging filed in B. n=94 ROIs. Scale bar, 50μm. (D) Example calcium traces (fluorescent value) of 4 selected neurons during the lever-press task. These neurons correspond to the neurons which color marked in C. The lever trajectory is shown in the top trace. Grey shading indicates movement epochs from successful trials.

### 2.2. Successful continuous decoding from layer 2/3 calcium signals

First of all, we tried to present the potential of these calcium signals from layer 2/3 in continuous neural decoding. For each neuron, calcium signals were calculated as the relative change in fluorescence, DF/F, over time. We classified these neurons as task-related or not using the Wilcoxon rank-sum test. 70.17% ± 10.83% of neurons in each session were task related. The results indicated the calcium signals from layer 2/3 neurons were indeed associated with forelimb movement. For a better understanding of neural content and coding, the RNN decoder with Hessian-Free (HF) optimization that allows information about past and future inputs to inform the predictions was used[19; 20]. We trained and test the HF-RNN decoder with 5-fold cross-validation. Only successful trials were included here (number of successful trials per session: 112 ± 16). The decoding performance was evaluated by Pearson’s correlation coefficient (CC) between predicted and actual pressure. All the results demonstrated high decoding performance (**Figure 2**). The average decoding performance for all sessions achieved up to 0.85 ± 0.06 (Figure 2A). The CC between the predicted and actual pressure was up to 0.93 for a single session (Figure 2B). The decoding performance from task-related neurons only was not inferior as well (Figure S2, Supporting Information). To prevent overfitting, we confirmed the generalization ability of trained decoders (Figure S3, Supporting Information).

**Figure 2.**
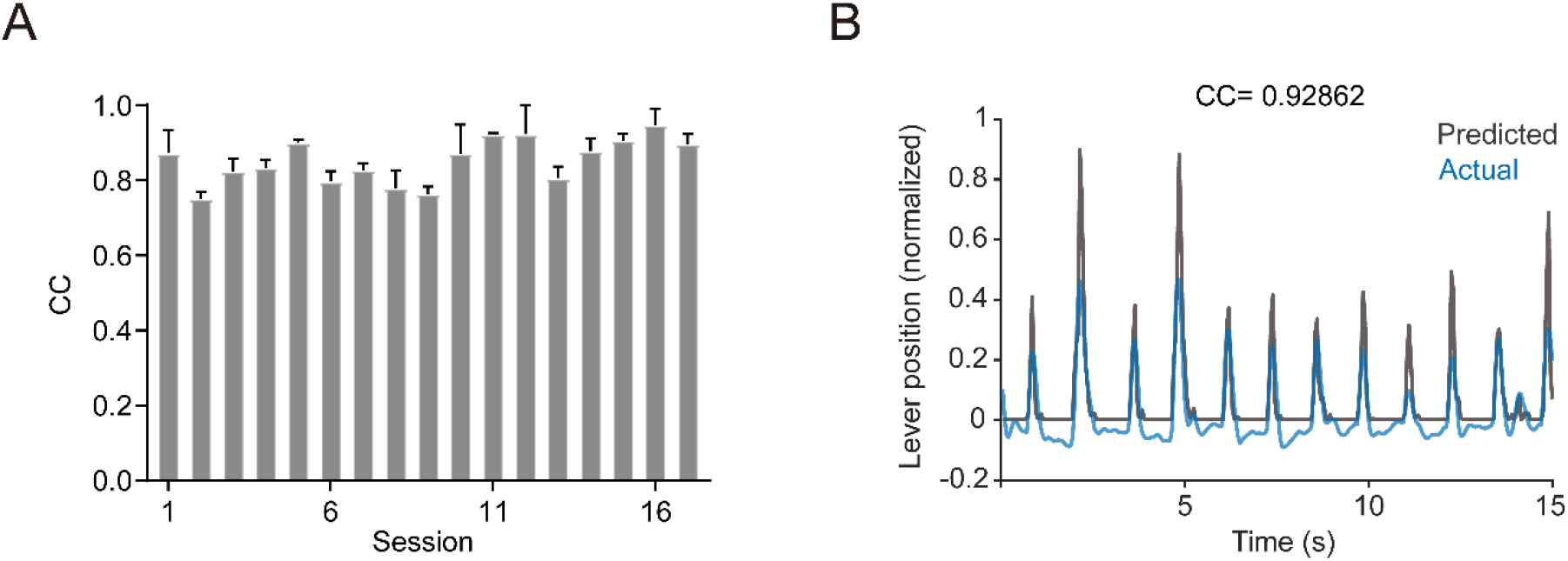
Continuous neural decoding using calcium signals from the layer 2/3 neurons. (A) Decoding performance through HF-RNN decoder in terms of CCs for all 17 sessions. Each performance was computed with 5-fold cross-validation. Each bar represents the mean ± SD. (B) Example of traces of the recorded lever trajectory (blue) and the lever trajectory predicted from the decoder (grey) in a single session.

### 2.3. The heterogeneous dynamics of layer 2/3 calcium signals

Considering successful continuous decoding from calcium signals, how the layer 2/3 neurons participated in movement coding was still unknown. Layer 2/3 neurons have been reported to integrate sensory and motor information from different input pathways. This different information suggested the existence of neurons with different motor representations. Wherefore we analyzed the activity patterns of these task-related neurons individually using trial-averaged DF/F traces. The calcium traces from all the successful trials in the session were aligned to movement onset and then averaged across the trials. The normalized resulting traces of task-related neurons were sorted based on their peak activation time and displayed in temporal raster plots (**Figure 3**A). We found that the neurons could be divided into three types according to their different temporal dynamics during the lever pressing, namely the pre-pressing, pressing, and post-pressing neurons. Their peak activation time was before pressing, up to pressing and after pressing respectively (Figure 3B). The remaining neurons that did not belong to these three types were defined as other task-related neurons.

**Figure 3.**
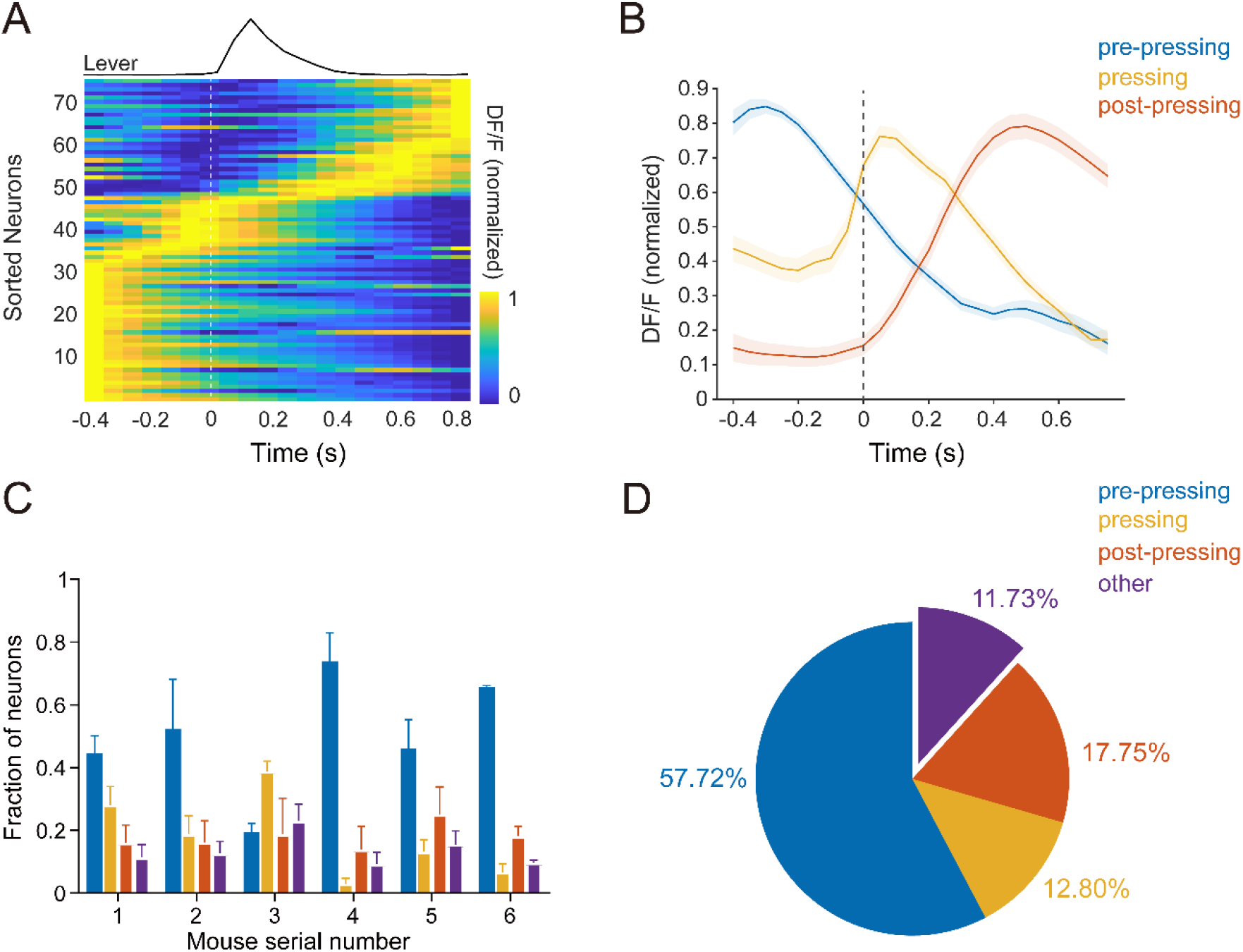
Dynamics of calcium activity of task-related neurons during the lever-press task. (A) Normalized average activity of all task-related neurons from mouse 5 in session 3 aligned to movement onset (dotted line). All the neurons were sorted based on their peak time during the task. n=77 neurons. (B) Representative trial-averaged calcium traces (DF/F±SEM) of pre-processing (blue), pressing (yellow) and post-pressing (red) neurons. The movement onset time is indicated by the black dotted line. (C) The quantity distribution of different types of task-related neurons across six mice. Error bars are SD. (D) The quantity distribution of different types of all task-related neurons, n=1594 (17 sessions from six mice).

Although the distribution of three main types of neurons was slightly different among six mice, the distribution across different sessions from the same mouse was relatively consistent (Figure 3C, Figure S4, Supporting Information). The pre-pressing neurons, as the dominant neurons, occupied 57.72% of all the task-related neurons. The percentage of pressing and post-pressing neurons was close, which was 12.80% and 17.75% respectively (Figure 3D).

However, we did not find obvious spatial clustering among the three types of activity patterns in 14 out of 17 sessions (Figure S5A, Table S1, Supporting Information), which suggested intermingled representations among different types of neurons in layer 2/3 of the motor cortex.

### 2.4. Different sparsity representations from different calcium dynamics

The three different calcium dynamics may play distinct roles in movement encoding. In the previous description, we noticed some neurons with a very sparse active pattern. We guessed whether the sparsity representations are associated with the specific calcium dynamics. As we all know sparseness is one of the principles important to sensory representations. Here we introduced kurtosis, referred to the definition of lifetime sparseness widespread in the sensory cortex, to measure the distribution of responses of neurons over time during the task-related period. The high kurtosis indicated a high degree of sparseness. We found that the pre-pressing and pressing neurons exhibited significantly higher lifetime sparseness compared to the post-pressing neurons, which was also consistent with the results displayed in Figure 2D (**Figure 4**A, Figure S6).

**Figure 4.**
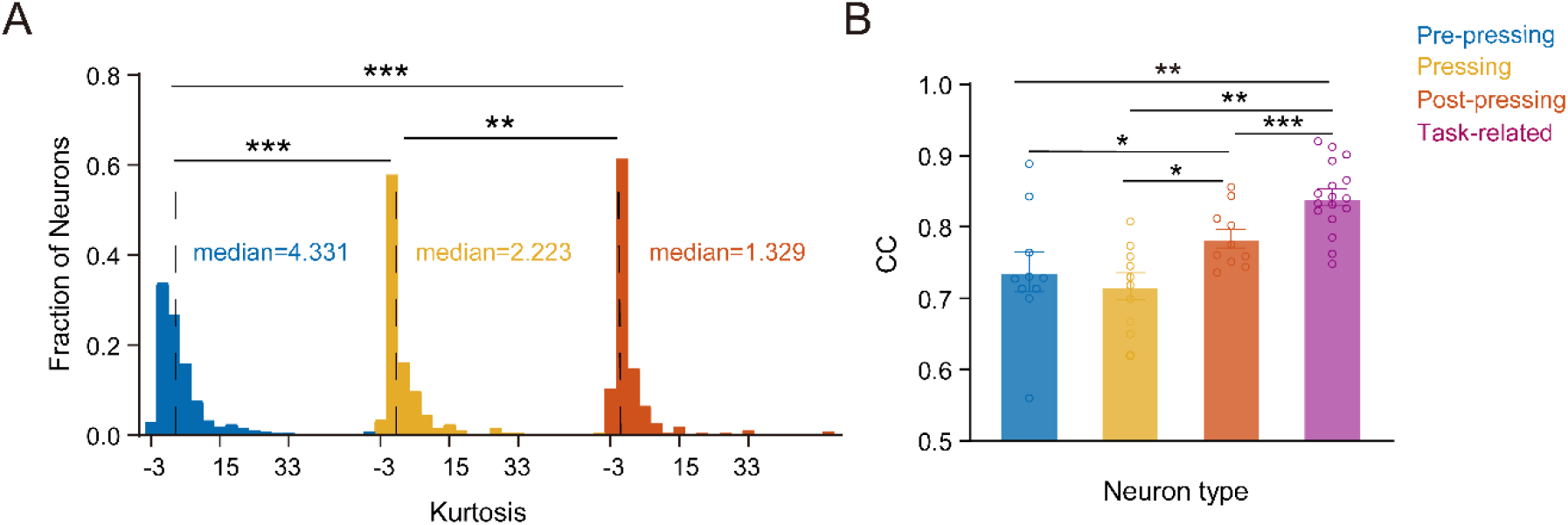
(A) The distribution of kurtosis measure to histograms of responses of an individual neuron overtime during the task-related periods. The median line is indicated by the black dotted line. ***p<0.001, **p<0.01, Kolmogorov-Smirnov test. (B) Comparing the decoding performance in terms of CC utilizing calcium signals from different neural ensembles. Error bar shows SEM. 10 sessions with at least 10 target neurons imaged are chosen for the three types of neurons. ***p<0.001, **p<0.01, *p<0.5, Wilcoxon rank sum test.

### 2.5. Consistency of population coding among different calcium dynamics

The question worth pondering is whether these different sparsity representations influenced population coding. Or the key to successful decoding was more from the post-pressing neurons with the lowest lifetime sparseness. To confirm this assumption, we compared the amount of movement information carried by these distinct populations. Not surprisingly, Post-pressing neurons encoded the lever trajectory most precisely among the three types of neurons. However, the pre-pressing neurons and pressing neurons with such sparse active patterns still showed good decoding performance with no significant differences. Certainly, none of the three types of neurons encoded the movement equally as the task-related neurons (Figure 4B). It can be seen that the three types of neurons are all involved in movement encoding at a considerable level, but the way they participated in population coding was different.

### 2.6. Different spatiotemporal coding patterns of different calcium dynamics

The comparable population decoding performance and the different degrees of sparseness hinted to us there may be distinct coding patterns from different calcium dynamics. We used the dimension reduction methods to characterize the population responses of the three types of neurons as well. The results displayed obvious spatial clustering in all 17 sessions, which proved their different coding patterns (Figure S7, Table S2, Supporting Information). We thence further explored the active properties of these calcium dynamics. We first investigated the relationships between the calcium dynamics and lever pressure individually. A large fraction of the three types of neurons all exhibited a low correlation to pressure value at the single neuron level. Even the highest CC was not more than 0.5 (**Figure 5**A). This demonstrated that the three types of neurons all encoded the pressure by the population responses rather than some individual functional neurons. At the same time, the mean CC values were ranked exactly the opposite of the kurtosis among the three types of neurons (Figure 5B, Figure 4A). The pre-pressing neurons with the highest lifetime sparseness had the lowest correlation. From this perspective, the sparseness of active patterns may partly explain the low correlation individually. In the previous description, the pre-pressing neurons and pressing neurons showed good decoding performance with no significant differences. Since there were no individual functional neurons, these sparse active patterns must complement each other for the movement representation. We calculated the correlation between the calcium dynamics itself among different types of neurons (Figure 5C, Figure 5D). The correlation between the post-pressing neurons was higher than the pre-pressing neurons and the pressing neurons. We also plot the correlation between the population responses and lever movement among different types of neurons (Figure 5E). As we expected, only pre-pressing and pressing neurons displayed the linear relationship between correlation and the number of neurons. The greater the number of neurons, the greater the correlation. We can see that the movement information carried by pre-pressing and pressing neurons was with higher complementarity. By contrast, the movement information from post-pressing neurons, which responded more regularly, was with higher redundancy.

**Figure 5.**
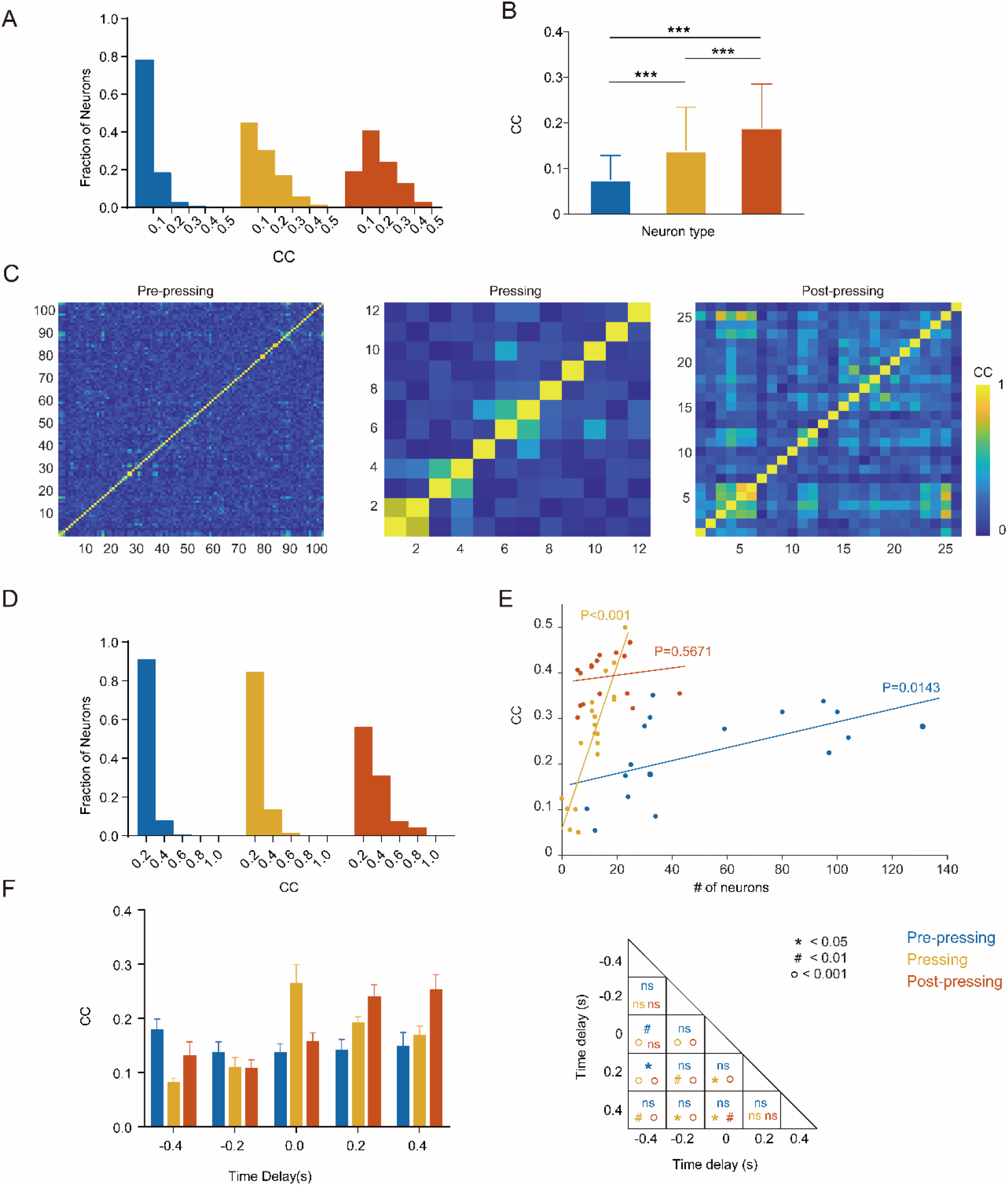
Characteristics of pre-pressing, pressing and post-pressing neurons. (A) Distribution of different types of neurons with different correlations to the pressure value. n=1408 neurons, 17 sessions from six mice (blue for pre-pressing, yellow for pressing, orange for post-pressing). (B) The mean correlation between the single neuron from different types and the pressure, n=1408 neurons, 17 sessions from six mice. ***p<0.001, Wilcoxon rank sum test. (C) The CCs between the calcium dynamics among the different types of neurons during one single session (left for pre-pressing, middle for pressing, right for post-pressing) (D) Distribution of CCs between the calcium dynamics among the different types of neurons. n=1408 neurons, 17 sessions from six mice. (E) The CCs between ensemble responses of different types of neurons and pressure (17 sessions from six mice). CC is plotted as a function of number of neurons. Each dot represents a single session. Regression lines with P values for the regression are shown for all points. (F) The decoding performance in terms of CC using Kalman decoder from pre-pressing (blue), pressing neurons (yellow), and post-pressing(red) neurons with different time delays. *p<0.05, #p<0.01, ο p<0.001, Paired-sample T test.

Besides that, although the pre-pressing neurons had the lowest individual correlation and highest lifetime sparseness, the dominant quantity made it comparable to pressing neurons at the population encoding (Figure 3D, 4B). However, the increase of CC caused by an increase in the quantity of pre-pressing neurons was much slower than that of post-pressing neurons (Figure 5E). The movement information conveyed by pre-pressing neurons was much less than the same quantity of post-pressing neurons. The coding patterns of post-pressing neurons made the decoding accuracy predictably better.

The three types of calcium dynamics displayed different temporal active patterns. The temporal dynamics itself may also affect the movement representation. Here we use a simple linear decoder, the Kalman decoder, to show the impacts of temporal dynamics on the movement representation. We performed time delay on the calcium signals and compared the decoding performance again. The activities of pre-pressing ensembles displayed the highest decoding performance when the time delay was -400ms, while the activities of post-pressing ensembles displayed the highest decoding performance when the time delay was 200ms and 400ms which better matched the press movement. Pressing ensembles predicted the movement most precisely without any time delay (Figure 5F). To some extent, we confirmed these calcium dynamics were related to lever-press movement with different temporal properties.

Taking together, although the three types of neurons all participated in lever movement coding through the population responses, the coding patterns were with distinct spatiotemporal characteristics.

## 3. Discussion

In this study, we used a miniscope to monitor somatic signals from hundreds of layer 2/3 neurons in M1 when mice performed a lever-press task. We not only demonstrated the great capability of somatic calcium signals from layer 2/3 in continuous neural decoding through HF-RNN but also further investigated how these layer 2/3 neurons participated in the movement coding. Our results provided some insights into the motor cortex study and the generation of optical BMIs.

We found three types of calcium dynamics with distinct spatiotemporal encoding patterns involved in the lever movement. In general, structure affects function. The multiple input pathways converging on M1 were likely to make the diverse motor representations here. In terms of the temporal active properties (Figure 3A, 3B), the calcium signals of pre-pressing neurons may be derived from the premotor cortex (M2). While the activity of post-pressing neurons was a response to sensorimotor cortex afferents or thalamocortical projections carrying sensory information[21]. We did not detect the obvious spatial clustering among the three types of patterns (Figure S5A, Supporting Information). Instead, they formed functional groups with predominantly local but also some long-range correlation (Figure S5B, Supporting Information). This is also consistent with the structure of intermingled representations of the motor cortex.

Different from some functional neurons serving as the good predictor of the movement in layer 5, the three types of calcium dynamics all exhibited low correlation to pressure at the individual level (Figure 5A). However, the movement information carried by these populations was capable of decoding lever pressure (Figure 4B). We can see that the specific kinematic parameters were only encoded by the ensembles in layer 2/3. One possible reason was the top-down control of the motor cortex, which led to more abstract information in layer 2/3 compared to layer 5[22; 23].

Although they all participated in movement encoding through the population. How they work was totally distinct. It’s noteworthy that we found the active patterns of pre-pressing and pressing neurons had a higher degree of sparseness (Figure 4A). It has been suggested that the most efficient neural codes perform sparse encoding[24]. There has been wide evidence of sparse coding in the sensory cortex especially in the visual cortex[25; 26]. While the sparse coding in the motor cortex was still controversial. Some evidence has been revealed in previous studies. Some motor neurons in layer 6 of the rabbit motor cortex will produce just one spike during some movement[27]. The stimulation of single neurons in the vibrissae motor cortex of rats can evoke whisker movements[28]. Our results here further support the presence of sparse coding in the motor cortex.

In the primate or rodent motor cortex layer 5, the activities in the preparation period were strongly related to the accurate movements that followed[29]. Even the pre-lever-movement peak was a better predictor of lever movement than neural activity recorded during the lever movement itself. In our study, these pre-pressing neurons, compared to pressing or post-pressing neurons, exhibited the lowest correlation to pressure value at the individual level (Figure 5A, Figure 5B). Meanwhile, at the population level, the post-pressing rather than pre-pressing neurons predicted the lever movement most precisely (Figure 4B). On the one hand, the difference in their coding patterns led to this difference. Pre-pressing neurons enabled sparse coding of lever trajectories through complementary spatiotemporal information. While Post-pressing neurons predicted the lever movement more accurately through the high fidelity and high redundancy of calcium dynamics.

On the other hand, the distinct roles played by layer 2/3 neurons and layer 5 neurons were also an important reason. The activities of layer 2/3 neurons intend to reflect the sensory input feedback and be a retrospective measure of movement performance itself. Levy S *et al*. reported the role of layer 2/3 pyramidal neurons in generating performance outcome-related signals during dexterous movements, which was a global assessment of motor performance rather than specific kinematic parameters or rewards. While layer 5 neurons encoded the specific parameters of behavioral response in the presence of sensory feedback to generate movement[30; 31]. Post-pressing neurons as the typical feedback of lever-press movement evoked or a retrospective measure of movement performance were more correlated to the lever-press movement. By contrast, the pre-pressing neurons here intended to work as a backup of the information from M2. During the movement preparation, such sparse and uncorrelated activity patterns of pre-pressing neurons were similar to independent basis functions, which provided more degrees of freedom for the following movement to generate a variety of pressure values. Therefore, such sparse active patterns are undoubtedly an optimal and efficient coding solution.

Despite the differences between layer 2/3 and layer 5 neurons, our results did present the capability of somatic calcium signals from layer 2/3 M1 in continuous neural decoding through high decoding performance, proving the feasibility of an optical motor BMI. Meanwhile, the movement information from the subpopulations was also adequate for specific kinematic parameters prediction. In an optical BMI system that processed image information, the time latency was specific to the recording location and the number of neurons recorded. A subpopulation of neurons recorded and processed enabled the implementation of low-time latency for a real-time optical BMI system.

One of the greatest superiorities of optical imaging technologies over electrophysiological methods is enabled advancements in genetics. In addition to excitation downstream from layer 2/3 to layer 5, in which layer 2/3 PNs contact the PNs in layer V. Layer 5 PNs contact inhibitory interneurons in layer 2/3[32]. Inhibitory and excitatory neurons in layer 2/3 during a forelimb lever-press task displayed different spatiotemporal activity[14]. The activity of inhibitory interneurons may also contribute to continuous decoding. A previous study conducted by Mitani *et al*. has presented the BMI with inhibitory interneurons. Even for inhibitory interneurons, different subtypes exhibited different neural dynamics[8]. The different types of calcium dynamics in our study may also come from distinct types of neurons or different projections. In the following studies, imaging for a specific cell type will help to uncover the function of distinct cell types in movement coding.

## 4. Methods

### 4.1. Subjects

All surgical and experimental procedures conformed to the Guide for The Care and Use of Laboratory Animals (China Ministry of Health) and were approved by the Animal Care Committee of Zhejiang University, China. Adult (8-12 weeks) male C57BL/6J mice were used for experiments. Mice were housed under standard housing conditions under a normal light cycle (12-h light/dark cycle) with food and water. Before the lever-press task training, mice were water restricted for at least 48 hours.

### 4.2. Behavior Task

Mice were first trained to perform a lever-press task similar to that previously described[33]. Each trial began with a 700ms, 500Hz tone. A successful trial was rewarded with water (10ul per trial) paired with a 350ms, 1000Hz tone, after which there was a variable inter-trial interval (ITI) lasting 3s-5s. A successful trial was defined as crossing two thresholds (a pressing threshold and a resetting threshold) and a holding period for at least 200ms in which the pressure exceeded the pressing threshold within response time (5s). The pressing threshold defined the pressure required for a successful lever press while the resetting threshold was used to prevent the mouse from holding the lever. Failure to pass the two thresholds during the response time or a timeout would both trigger a 150ms, 100Hz tone (Figure S1C, Supporting Information). A lever press during ITI was neither rewarded nor punished. Each session consisted of 100-200 trials that lasted 20-30 minutes. When the mice accomplished 120 trials or they stopped performing tasks, a session would terminate. Mice were trained with this task daily for 10 days before surgery.

### 4.3. Surgery

Mice (n=6) that successfully learned this task after a 10-day behavioral training were anesthetized with isoflurane and intraperitoneally injected with ketamine (100 mg/kg) and xylazine (10 mg/kg). Then a midline incision of the scalp was made to expose the periosteum, and the skull above the left caudal forelimb area (CFA) was located based on stereotactic coordinates (from -1.5 to +1.0 mm of bregma and from 0.5 to 3.0 mm lateral to midline; Allen Mouse Brain Atlas) and marked with ink. A craniotomy (3 mm diameter) was performed over the left CFA. Adeno-associated viruses (AAV) carrying genes for the calcium indicator jGCaMP7f (pGP-AAV-syn-jGCaMP7f-WPRE, Obio Technology Corp) were injected into the left CFA of the motor cortex (bregma +0.3 mm, lateral 1.5 mm, depth 0.25 mm). After viral injection, a glass coverslip (3.0 mm diameter) was glued to the skull above the exposed dura (Figure S1A, Supporting Information). The scalp was sutured to protect the surface of the glass coverslip. Three weeks later, the expression of jGCaMP7f was appropriate and a custom baseplate was glued over the glass coverslip with an optimized focus to observe the neuronal activity. Mice were then sent to recover from the surgery for 1 week. Before imaging, mice with head baseplates were habituated to the imaging apparatus three times (10 min each) to minimize the potential stress effects of head restraining and imaging. No obvious distress was observed in habituated animals during imaging experiments.

### 4.4. Data Acquisition

Calcium signals from layer 2/3 CFA were imaged via UCLA miniscope, a one-photon imaging system. Imaging data were acquired at 752×480 pixels with a frame rate of 20Hz in a field of view of 700×450 μm. The depth of imaging was approximately 150-250 μm from the dura for Layer 2/3 (Figure S1B, Supporting Information). The time of each session was kept to less than 30 minutes to prevent photonic bleaching. Figure S1C shows the overall experimental pipeline for the lever-press task including a combination of calcium imaging and behavior setup (Figure S1C, Supporting Information). The behavior setup was conducted by visual studio software, which controlled task events including task cues, threshold crossing monitoring and reward control. During the trials, the voltage from the force transducer sampled at 30 kHz, which is proportional to the lever pressure, was continuously recorded by neural simultaneous processing (NSP), a Cerebus acquisition system. Except for pressure, the timestamps for stream alignment from the miniscope and behavior setup were also synchronously recorded by NSP.

### 4.5. Movement analysis

During the lever-press task, movement data was identified in the lever pressure (voltage recordings from the force transducer). The data was firstly baseline removed and then filtered with a 2-order, 20 Hz low-pass Butterworth filter. After that, timestamps from the miniscope were used to downsample the data from 30k Hz to 20 Hz. The movement onset time was defined by finding when the lever pressure crossed a threshold exceeding the resting period before the movement, and the end time was defined by finding when the lever pressure went below the resetting threshold following the movement[14].

### 4.6. Processing of calcium imaging videos

All analyses were performed using MATLAB (Mathworks). After getting the raw videos, we performed neural enhancement, movement correction and neural signal extraction with a miniscope 1-photon-based calcium imaging signal extraction pipeline (MIN1PIPE) developed by Jinghao Lu *et al*. [18]. Thus, ROIs with corresponding calcium signals would be extracted. The resolution was approximately 0.93μm/pixel for the UCLA miniscope. We set the size of the structure element comparable to half of the overall size of the neurons as 9 in MIN1PIPE parameter settings for ROIs identification, which matched the size of soma instead of the dendritic trunk.

#### 4.6.1. Fluorescence analysis

For each ROI, calcium signals were calculated as the relative change in fluorescence, DF/F= (F-F0)/F0, over time. The F0 was estimated as the baseline of fluorescence from a ± 15 s sliding window. To visualize the activity patterns of the single neuron during the lever-press task, we calculated trial-averaged DF/F traces for each ROI. The DF/F value in each trial was calculated from 400ms before the movement onset to the end time of movement, which is defined as the task-related epochs. The DF/F traces of each ROI were aligned by movement onset and then averaged across trials. Only successful trials were included. The resulting traces from all ROIs were sorted based on their peak time during the task and displayed in temporal raster plots.

#### 4.6.2. Classification of different types of calcium dynamics

The neurons were classified into different types according to the method in a previous study [17]. Briefly, among all identified neurons, an individual neuron was defined as task-related when DF/F during all task-related periods was significantly larger (p<0.05 in Wilcoxon rank-sum test) than DF/F during all ITIs. For each task-related neuron, DF/F was then aligned to either the onset or end time of all successful lever movements at time 0 and then averaged over all successful trials and defined as *F*_*start*_ and *F*_*end*_, respectively. The neurons with peak activity 20% larger than mean activity during task-related periods were collected. Neurons with *F*_*start*_ at peak times of < 0s were considered pre-pressing neurons. For neurons with *F*_*start*_ at peak times of > 0s, the peak amplitude before the end of the lever press, *max*_*t*≤0_*F*_*end*_(*t*), was compared with the peak amplitude after the end of the lever pressing, subtracted by the exponential decay component of the amplitude at the end of the lever pressing, *max*_*t*>0_(*F*_*end*_(*t*)− *F*_*end*_(0)*e*^−*t*/1^). Neurons were classified as pressing neurons when *max*_*t*≤0_*F*_*end*_(*t*) was larger than *max*_*t*>0_(*F*_*end*_(*t*)− *F*_*end*_(0)*e*^−*t*/1^). The remaining neurons were considered post-pressing neurons. Task-related neurons that did not belong to these three types were defined as other task-related neurons.

#### 4.6.3. spatial clustering analysis

The spatial clustering index (SCI) is used to quantify spatial clustering. SCI is the ratio of the mean distance between neuronal pairs belonging to different classifications divided by the mean distance between neuronal pairs belonging to the same classifications exclusively. Under the null hypothesis that different classifications are intermingled, SCI ∼ 1. SCI > 1 indicates spatial clustering. We shuffled the cell’s labels and computed SCI on the shuffled data 1000 times to test for statistical significance. The p-value is the fraction of shuffled SCI higher than the unshuffled SCI.

#### 4.6.4. Sparseness measure

The sparseness measure is used to quantify the peakedness of the response distribution. Two definitions of sparseness are widespread. One is to describe codes in which few neurons are active at any time (‘population sparseness’), and the other is to describe codes in which each neuron’s lifetime response distribution has high kurtosis (‘lifetime sparseness’) [34]. We used the latter in our study. The calcium signals of each neuron during task-related periods from all successful trials were used only. The lifetime sparseness uses kurtosis (the fourth statistical moment of distribution) as its metric. The equation is as follows:

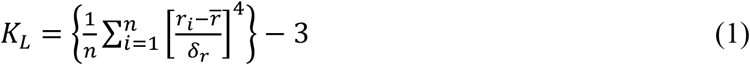

where 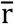 and δ_*r*_ are the mean and standard deviation of the responses. A distribution with high positive kurtosis means it contains many responses which are small (compared to the δ_*r*_), and only a few responses that are very large. A distribution will have large negative kurtosis if all responses are present equally often in the distribution. The Gaussian distribution has the zero kurtosis.

#### 4.6.5. Dimension reduction and visualization

Dimension reduction methods, which are well-suited for analyzing neural population activity, were used to produce low-dimensional projections from high-dimensional data and realize data visualization[35]. Considering nonlinear manifold discovery, we applied a nonlinear t-distributed stochastic neighbor embedding (t-SNE) algorithm here. t-SNE can capture much of the local structure of the high-dimensional data, while also revealing global structures such as the presence of clusters at several scales[36]. In order to reduce computational complexity, the high-dimensional data was first projected into 10 decor-related components by principal component analysis (PCA). The t-SNE then refined the PCA results to produce a 2-dimensional mapping[37].

### 4.7. Continuous Decoding

For continuous decoding, we adopted an improved recurrent neural network (RNN) decoder. The biggest difference between this decoder and standard RNN is that the training used Hessian-Free optimization (HF) together with a novel damping scheme [19; 20]. We used the DF/F traces from all or subsets of neurons during the task-related epochs and ITIs from all successful trials to train and test the HF-RNN decoders with 5-fold cross-validation. For the generalization ability test of trained decoders, the calcium signals across the entire trials were used as the test data. The decoding performance was evaluated by Pearson’s correlation coefficient (CC) between predicted and actual pressure.

### 4.8. Quantification and Statistical analysis

Typically, n=6 mice were used for analyses of the results in the lever-press task. Statistical analyses are described in the results, and figure legends. In general, the paired t-test or Wilcoxon rank sum test was used to determine the difference between groups. P < 0.05 was considered statistically significant.

### 4.9. Histology

Mice were deeply anesthetized with isoflurane and then perfused transcardially with PBS followed by 4% paraformaldehyde. The brain was removed and postfixed with 4% paraformaldehyde overnight at 4 °C. After that, the brain was dehydrated with 30% sucrose in PBS solution over 48 hours at 4 °C. After freezing, the brain was sectioned into 50μm coronal slices. Slices were imaged under a fluorescence microscope (Olympus) with a 10×and 20×objective and a fluorescence filter was set appropriately for GFP. The expression of jGCaMP7f in the motor cortex of all mice used for in vivo imaging was confirmed (Figure 1B). Selected slices were imaged using a confocal microscope (Zeiss).

## Author Contributions

Ruixue Wang and Jiawei Han conceived and performed the experiments. Ruixue Wang participated in data analysis and wrote the manuscript with input and editing from all authors. Shaomin Zhang provided guidance and assistance with all aspects of the work.

## Data Availability

The datasets and code for this study are available from the corresponding authors on reasonable request.

