## Supporting information for "Somatic Calcium Signals from Layer II/III Motor Cortex for Continuous Neural Decoding"

### 1. Supplementary Figures

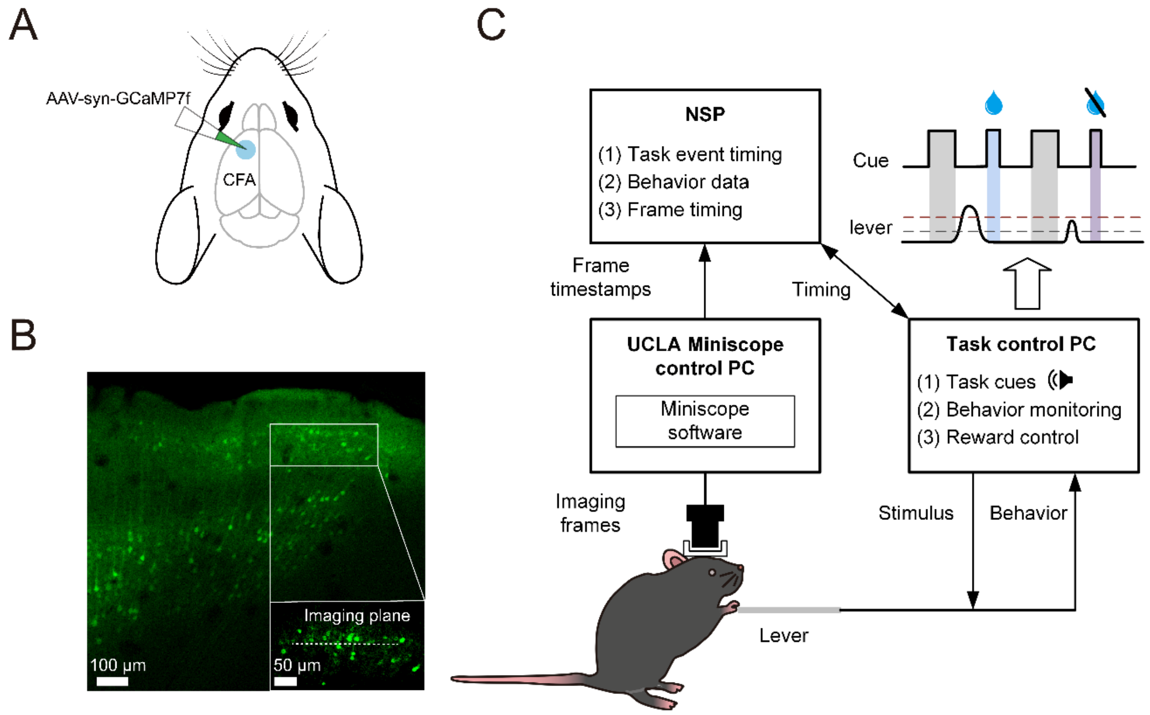


**Supplementary Figure 1.** Lever-press task under the miniscope. (A) Diagrams showing GCaMP7f labeling. (B) A representative image showing the Gcamp7f expression in layer 2/3 motor cortex. Dotted line: imaging plane. Scale bars: overview, 100μm; inset, 50μm. (C) Experimental pipeline for synchronous recording of one-photon imaging data and behavior data. NSP: neural simultaneous processing. Right top: Schematic of the event timing for each trial. Shading indicates different cues (grey for start, blue for reward, purple for failure). Black dotted line: resetting threshold. Red dotted line: pressing threshold.

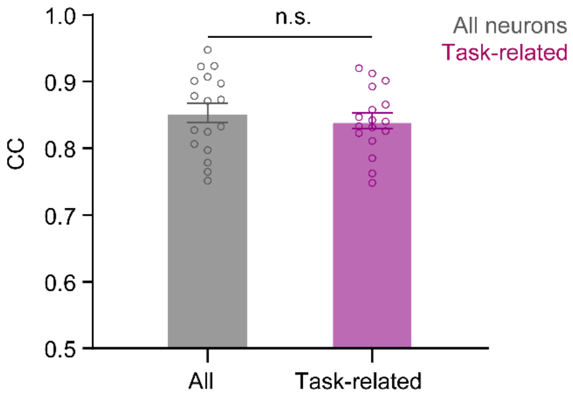


**Supplementary Figure 2.** Comparing the decoding performance in terms of CC utilizing calcium signals from all neurons and task-related neurons (17 sessions from six mice). Each performance was computed with 5-fold cross-validation.Error bar shows SEM. **p<0.01, Paired-sample T-test.

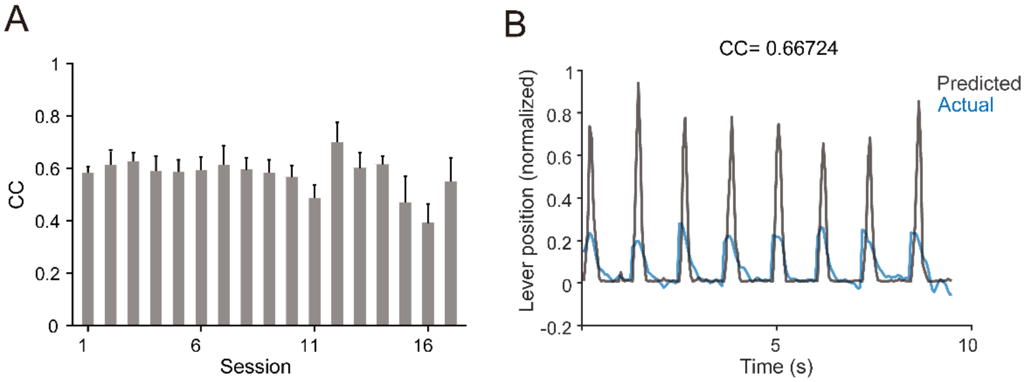


**Supplementary Figure 3.** The Continuous neural decoding using calcium signals from all layer 2/3 neurons. (A) Generalization ability presented by decoding performance through HF-RNN decoder in terms of CCs for all 17 sessions. Each performance was computed with 5-fold cross-validation. Each bar represents the mean ± SD. The mean CC for all sessions was approximately 0.6 (B) Example of traces of the recorded lever trajectory (blue) and the lever trajectory predicted from the decoder (grey) in a single session.

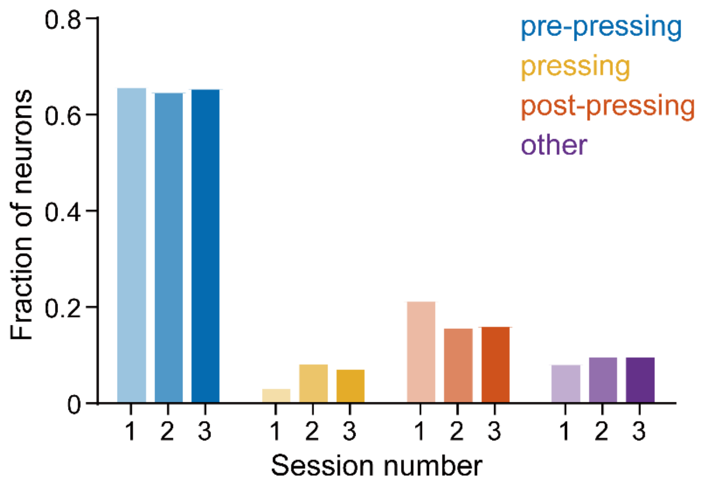


**Supplementary Figure 4.** An example distribution of different types of task-related neurons from mice 6. The distribution across different sessions from the same mouse was relatively consistent.

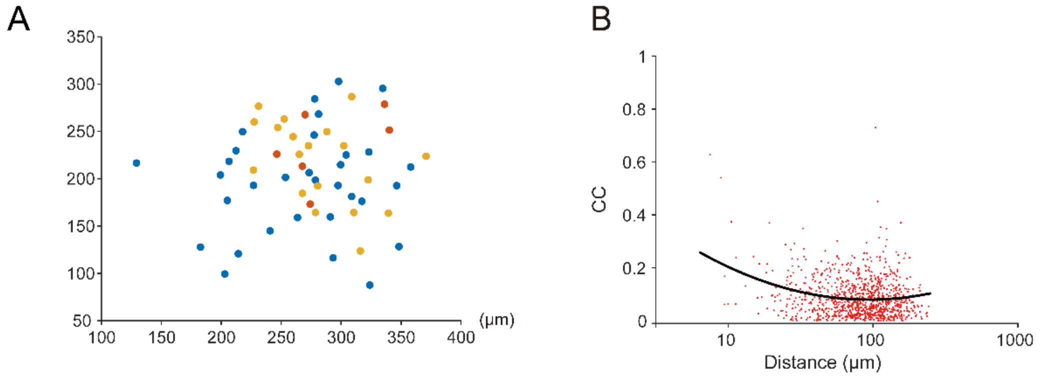


**Supplementary Figure 5.** (A) An Example image from mouse 1 showing the spatial mixed distribution of three types of neurons, n=63 neurons. SCI with significance was used to quantify the spatial clustering across different types of neurons. SCI=0.95, P>0.05. SCI > 1 with p<0.05 indicates significant spatial clustering. (blue for pre-pressing, yellow for pressing, red for post-pressing). The SCI with p-value for each imaging field is listed in the following Supplementary Table 1. (B) The relative position of neuronal pairs and corresponding CCs between DF/F traces during the movement epochs for a single session. CCs are plotted as a function of pairwise neuronal distance. Each dot corresponds to one neuronal pair. Red: individual pairs. Black: average. For better visualization, the distance axis was logarithmically scaled. The ensemble activity showed, on average, increased CCs for nearby neurons, but we also found some neurons even far from each other could exhibit relatively high CCs.

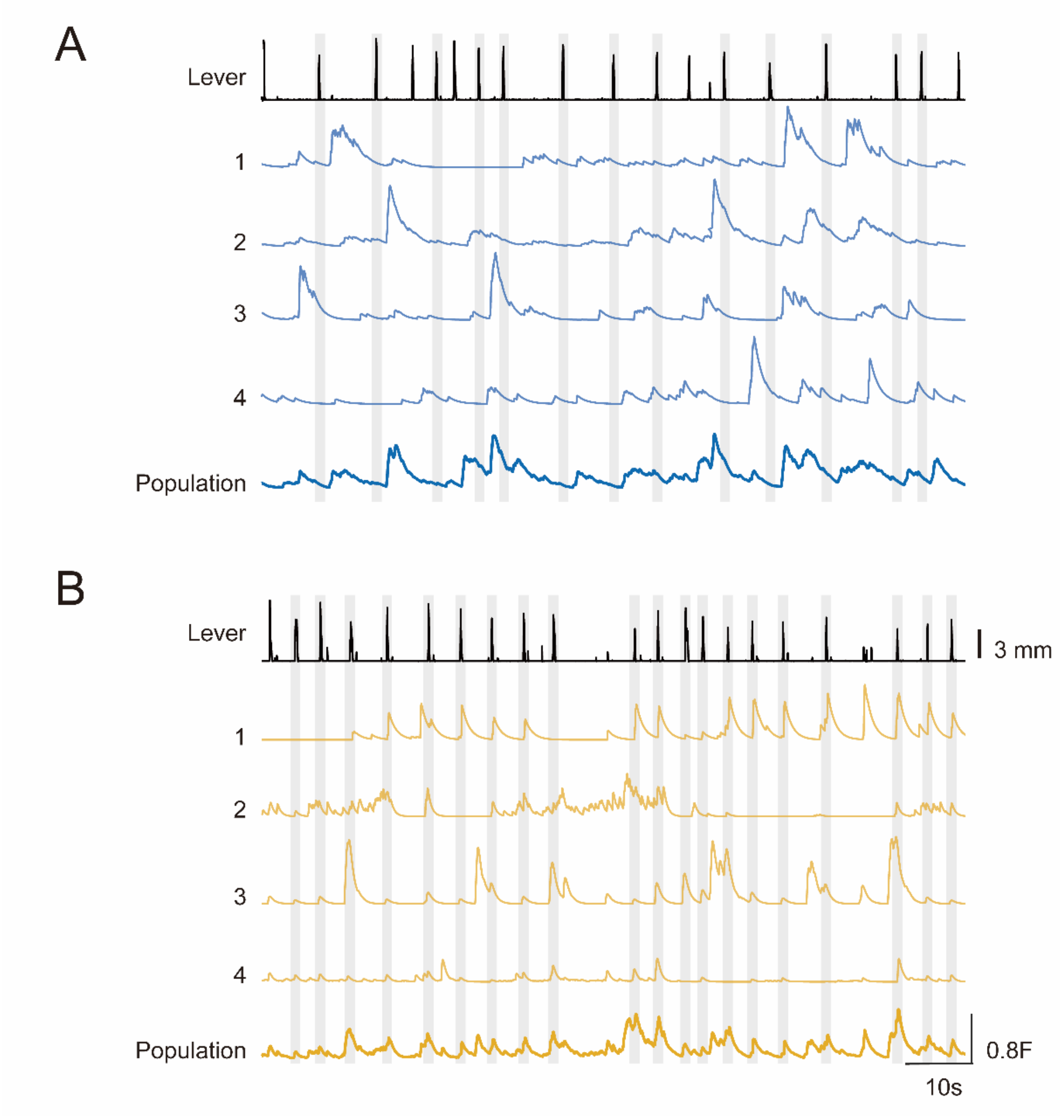


**Supplementary Figure 6.** Example calcium traces (fluorescent value) of (A) pre-pressing neurons and (B) pressing neurons during the lever-press task. Top: the lever trajectory. The calcium traces of the single neuron are randomly selected. The population response is shown in the bottom trace. Grey shading indicates movement epochs from successful trials.

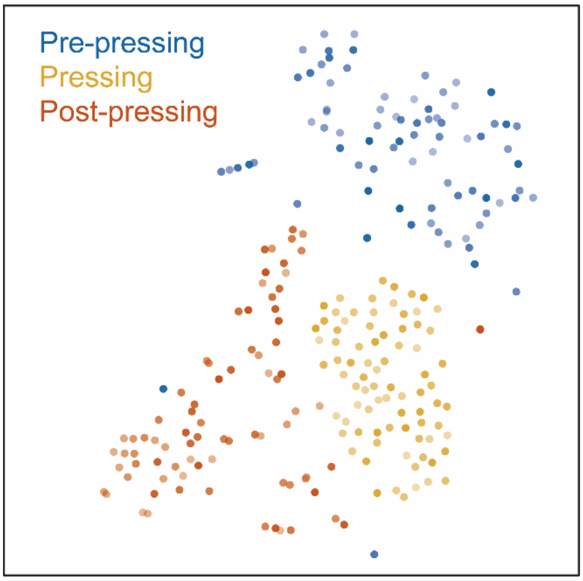


**Supplementary Figure 7.** Dimension Reduction and visualization for activities of pre-pressing(blue), pressing(green) and post-pressing(orange) ensembles in a single session. Each dot represents a trial. Trial number, n=99. The shade of color was mapped to the lever pressure. The darker color corresponds to the larger pressure. The spatial clustering index (SCI) with significance was used to quantify the spatial clustering across different ensemble activities. SCI=2.1611, p<0.001.

### 2. Supplementary Tables

**Supplementary Table 1.** The spatial clustering of the three main type neurons in the imaging fields

| Session | Spatial cluster index | P value |
|---|---|---|
| 1 | 0.9902 | 0.584 |
| 2 | **1.0886** | 0.037* |
| 3 | 0.9516 | 0.899 |
| 4 | 1.0116 | 0.366 |
| 5 | 1.0497 | 0.07 |
| 6 | 0.9914 | 0.537 |
| 7 | 1.005 | 0.403 |
| 8 | 0.9598 | 0.885 |
| 9 | 1.0456 | 0.148 |
| 10 | **1.2109** | 0*** |
| 11 | **1.17** | 0.008** |
| 12 | 1.0158 | 0.337 |
| 13 | 0.9454 | 0.77 |
| 14 | 0.94 | 0.9 |
| 15 | 0.99 | 0.57 |
| 16 | 0.9313 | 0.95 |
| 17 | 1.0412 | 0.12 |

SCI > 1 with $p<0.05$ indicates significant spatial clustering. *$p<0.05$, **$p<0.01$, ***$p<0.001$. There was no significant spatial clustering among the three types of calcium dynamics in 14 out of 17 sessions.

**Supplementary Table 2.** The spatial clustering the three ensemble activities in the low-dimensional representations

| Session | Spatial cluster index (SCI) | P value |
|---|---|---|
| 1 | **1.28** | 0*** |
| 2 | **1.4081** | 0*** |
| 3 | **1.8613** | 0*** |
| 4 | **1.2993** | 0*** |
| 5 | **1.383** | 0*** |
| 6 | **1.391** | 0*** |
| 7 | **1.4896** | 0*** |

| | | |
|---|---|---|
| 8 | **1.4598** | 0*** |
| 9 | **2.2367** | 0*** |
| 10 | **2.0086** | 0*** |
| 11 | **2.8574** | 0*** |
| 12 | **2.1611** | 0*** |
| 13 | **1.1723** | 0*** |
| 14 | **2.2663** | 0*** |
| 15 | **1.5039** | 0*** |
| 16 | **1.4707** | 0*** |
| 17 | **1.2809** | 0*** |

SCI > 1 with $p<0.05$ indicates significant spatial clustering. ***$p<0.001$. There was significant spatial clustering among the three ensemble activities in the low-dimensional representations in all 17 sessions.
